# Species-specific versus community-wide assays in eDNA monitoring of the European eel *Anguilla anguilla*: Trade-offs between detection sensitivity and the value of additional community data

**DOI:** 10.64898/2026.03.19.712641

**Authors:** Angus I.T. Monaghan, Graham S. Sellers, Nathan P. Griffiths, Lori Lawson Handley, Bernd Hänfling, James A. Macarthur, Rosalind M. Wright, Jonathan D. Bolland

## Abstract

Effective monitoring of the critically endangered European eel (*Anguilla anguilla*) is essential for conservation planning and regulatory decision-making, particularly in heavily fragmented rivers. Environmental DNA (eDNA) methods offer sensitive alternatives to traditional surveys, but there is uncertainty around whether targeted assays or community-wide approaches are better suited to achieve monitoring objectives. We compared eDNA metabarcoding and species-specific quantitative PCR (qPCR) for detecting *A. anguilla* across 145 pumped catchments in the Fens, East Anglia, England. All sites were sampled once initially, and sites negative for *A. anguilla* were re-sampled based on metabarcoding results. This allowed comparison of detection rates from a single water sample and site-level retrospective identification of sites where qPCR could have identified *A. anguilla* in earlier samples. The findings were also set in the context of the wider biodiversity information generated by metabarcoding. From the initial (single) water sample, qPCR detected *A. anguilla* at seven more sites than metabarcoding (17 versus 10). With repeated sampling, metabarcoding detected *A. anguilla* at 43 sites, including all but one of the sites where qPCR detected *A. anguilla*, and ten sites where qPCR did not detect *A. anguilla* within the same number of samples. Indeed, the additional sampling effort required to detect *A. anguilla* with metabarcoding at sites also positive with qPCR was small relative to the overall sampling effort. Furthermore, metabarcoding additionally detected 28 non-target fish species alongside fish, amphibian and mammal species of conservation concern. Our results highlight trade-offs between target-species sensitivity and the broader ecological information provided by each method, and support metabarcoding as an effective tool for a holistic conservation approach, with the additional community data outweighing the marginally increased sensitivity of qPCR.

## Introduction

Routine biodiversity monitoring plays a key role in ecological assessment and the identification of emerging conservation issues (Nichols & Williams, 2006). Assessment of distributions is particularly important for diadromous fishes (Ouellet et al., 2022) given their susceptibility to habitat loss (Verhelst et al., 2021). Analysis of traces of DNA in environmental samples (environmental DNA or ‘eDNA’) to determine species distributions has become a widely utilised tool for biodiversity monitoring in aquatic ecosystems, particularly for the detection of rare species (Jerde et al., 2011; Harper et al., 2018, 2019; White et al., 2020). Two of the main eDNA approaches are metabarcoding and quantitative PCR (referred to hereafter as qPCR). Metabarcoding uses polymerase chain reaction (PCR) with conserved primers to amplify entire biological communities, followed by high-throughput sequencing to detect a wide range of freshwater fish, vertebrates (Riaz et al., 2011; Miya et al., 2015) and other taxa (Leese et al., 2021; Cowgill et al., 2025). When compared with traditional methods, metabarcoding consistently detects as many and often more freshwater fish species (Hänfling et al., 2016; Li et al., 2019; Griffiths et al., 2020). By contrast, qPCR targets a specific taxon of interest, and is often more sensitive than traditional methods when targeting freshwater fish (Jerde et al., 2011; Wilcox et al., 2016; Weldon et al., 2020; Penaluna et al., 2021).

Choosing which eDNA analysis method to use when trying to establish the distribution of a target species in an unstudied location is a conundrum many researchers and regulators face. Metabarcoding is often favoured when assessing entire communities or several key species (Hänfling et al., 2016; Holman et al., 2019; Lawson Handley et al., 2019; Li et al., 2019; Mychek-Londer et al., 2020), while qPCR is often favoured when assessing distribution of rare or invasive or conservation priority species (Harper et al., 2018; Peixoto et al., 2023). Studies comparing qPCR and metabarcoding consistently show qPCR to have a higher degree of sensitivity to the target taxa DNA than metabarcoding (Harper et al., 2018; Blackman et al., 2020; Chandelier et al., 2021; Pont et al., 2022; Yu et al., 2022). qPCR is also faster and marginally more cost-effective than metabarcoding when targeting a single species (Harper et al., 2018). However, no studies explicitly consider qPCR sensitivity relative to the amount of additional metabarcoding effort to detect a target species at a location (i.e. how much sampling effort could qPCR save compared to metabarcoding when searching for a target species), nor do they compare it against the benefits of wider community composition data generated.

The European eel (*Anguilla anguilla*) is a ‘Critically Endangered’ (Pike et al., 2020) fish species with a catadromous lifecycle (van Ginneken & Maes, 2005), which leads to particular and challenging requirements for effective conservation (Knights et al., 2009; Buysse et al., 2014; Kerr et al., 2015; Verhelst et al., 2021). Land drainage and flood-relief infrastructure, including pumping stations, are particularly problematic for *A. anguilla*. The catchments upstream of pumping stations are sometimes suitable for juvenile *A. anguilla* to live and mature in (Laffaille et al., 2004) but upstream passage is obstructed or prevented, despite glass eel climbing abilities (Podgorniak et al., 2016). Downstream passage through pumps (entrainment) is a significant issue for mature ‘silver’ eels attempting to return to the Sargasso Sea to spawn (Buysse et al., 2014; Bolland et al., 2019; van Keeken et al., 2021; Evans et al., 2024). To conserve *A. anguilla* most effectively in pumped river systems, as per regulatory requirements (Council Regulation (EC) No 1100/2007, 2007; The Eels (England and Wales) Regulations 2009), a thorough understanding of their present-day distribution and habitat requirements is essential (Harwood et al., 2022). Both eDNA metabarcoding (Griffiths et al., 2020) and qPCR (Weldon et al., 2020) have demonstrated higher detection rates for *A. anguilla* than those of traditional fish sampling methods. However, while there are numerous studies comparing eDNA metabarcoding and qPCR, there are no such studies specific to *A. anguilla*.

Rivers upstream of pumping stations are rarely surveyed and thus there is very limited information on the presence/absence of *A. anguilla* and other aquatic species. Understanding the composition of the entire fish community is important at these sites given resident fish species are also vulnerable to entrainment (Norman et al., 2023a, 2023b) and can be used as indicators of habitat quality/connectivity (Blabolil et al., 2017; Sun et al., 2022), and thus likelihood of *A. anguilla* presence (Griffiths et al., 2025a). The distributions of conservation priority and non-native fish, amphibian and mammal species, given implications for management, would also be advantageous. For example, spined loach (*Cobitis taenia*), a species protected under European Commission legislation are well-known to inhabit drainage systems and have challenging monitoring requirements (Nunn et al., 2014). This is also true for water vole (*Arvicola amphibius*), a priority conservation species in the UK that are protected by the Wildlife and Countryside Act of 1982 (Richards et al., 2014). Notwithstanding, gathering knowledge of the entire fish, amphibian and mammal community cannot be at the expense of highly accurate assessments of target species’ distribution. Therefore, while eDNA metabarcoding would be the preferred sampling method, given the ability to detect both *A. anguilla* and the entire fish and wider vertebrate community, qPCR sampling may provide a more accurate assessment of *A. anguilla* distribution.

The study aimed to compare eDNA metabarcoding and qPCR as detection methods for *A. anguilla* across pumped catchments in East Anglia, England. The qPCR analysis was retrospectively performed on two targeted sub-sets of samples after all water sampling and metabarcoding sequencing had been completed with three objectives. Specifically, the first objective was to compare sample-level *A. anguilla* detection rate and agreement between the two eDNA methods for (i) the initial (single) water sample taken from all pumped catchments, and (ii) all samples from *A. anguilla*-positive sites using metabarcoding (at any point during the study). The second objective was to compare site-level *A. anguilla* detection rate agreement between the two eDNA methods and to quantify the number of additional water samples required to detect *A. anguilla* with metabarcoding rather than qPCR, referred to here as excess metabarcoding effort. The third objective was to compare the benefits of non-*A. anguilla* species detected with metabarcoding, particularly conservation and non-native species, and associations with *A. anguilla*. The findings will help inform debates surrounding which eDNA sampling method to use when attempting to establish the distribution of target species, potentially at a large spatial scale, to inform legislative/conservation action and infrastructure modification.

## Materials and methods

### Study sites

Water samples were collected from 145 pumping station catchments in the East Anglia Fenland region, England. The catchments are all part of low-lying artificial drainage systems in predominantly agricultural land, draining into the River Great Ouse and River Nene. Site access was granted by the Environment Agency and relevant Internal Drainage Boards; no specific permits or licences were required for water sampling at the locations studied. The “removal” survey design, as described by Mackenzie & Royle (2005) was used; only sites negative for the target species (*A. anguilla*) were subsequently re-sampled to maximise confidence in *A. anguilla* presence/absence, while minimising time and consumable costs. Water was sampled on four occasions or rounds; a single water sample was collected in May 2021 (R1), and two water samples were collected in March (R2) and July (R3) 2022, and September 2023 (R4). *A. anguilla*-positive sites with more than 100 eDNA sequence reads from metabarcoding were not re-sampled. Eighteen sites deemed very poor habitat quality and highly unsuitable for *A. anguilla* were not re-sampled after R1. In addition, 26 *A. anguilla*-negative sites that were inhibited in May 2021 (R1) had an additional sample taken in September 2023 (R4).

### Sample collection

Sampling equipment (sampling pole, cool box and sample bottles) were sterilised using 10% bleach solution at each site. Two-litre surface water samples were all collected directly upstream of each pumping station’s weed screen. Field negatives (1 per day of sampling) consisting of purified, deionised water were used each day. Samples were kept on ice in the field and refrigerated overnight. The following day, samples were filtered using vacuum pumps and sterile 0.45 μm mixed cellulose ester membrane filters (2 per sample) following the methodology described in Griffiths et al. (2025b). Filters were frozen at -20 °C in sterile 5 ml tubes.

### DNA Extraction

DNA extraction followed the “Mu-DNA” water protocol published by (Sellers et al., 2018) and also followed by Griffiths *et al*. (2025b). After the first sampling round, an additional ethanol wash step was introduced to for all samples to minimise inhibition. Extraction negatives (one per twelve of extractions) were used.

### Metabarcoding

Library preparation was carried out following the workflow detailed in Griffiths *et al*. (2025b), a summary follows. Nested metabarcoding, utilising a two-step PCR protocol, was carried out using multiplex identification (MID) tags in each step to allow sample identification, described by (Kitson et al., 2019). The first PCR (PCR1) was performed in triplicate (three PCR replicates per extracted DNA sample), amplifying a 106 bp fragment using published 12S ribosomal RNA primers 12S-V5-F (′5-ACTGGGATTAGATACCCC-3′) and 12S-V5-R (5′-TAGAACAGGCTCCTCTAG-3′) (Riaz et al., 2011; Kelly et al., 2014). Negative PCR controls (Molecular Grade Water, one per 24 reactions) were used throughout, alongside positive controls of DNA (0.05 ng μL^−1^) from non-native cichlids: *Maylandia zebra* and *Astatotilapia calliptera*. All PCR replicates were pooled after amplification, and samples from each PCR1 batch were normalised and pooled into sub-libraries which were then purified with MagBIND RxnPure Plus magnetic beads (Omega Bio-tek Inc., Norcross, GA, USA), using a double size selection protocol (Quail et al., 2009). Ratios of 0.9× and 0.15× beads to 100 μL of product from all sub-libraries were used. Next, the cleaned product underwent a second shuttle PCR (PCR2) to bind Illumina adapters to all sub-libraries. A second purification was performed on the PCR2 product with Mag-BIND RxnPure Plus magnetic beads (Omega Bio-tek Inc., Norcross, GA, USA). Ratios of 0.7× and 0.15× magnetic beads to 50 μL of each sub-library were used. Cleaned sub-libraries were stored at 4 °C until quantification and normalisation. After pooling, the final library was purified once more (following an identical protocol to the second clean), quantified by RTPCR using the NEB Next Library Illumina Quant Kit (New England Biolabs Inc., Ipswich, MA, USA). Fragment size and purity were then verified using an Agilent 2200 TapeStation using High Sensitivity D1000 ScreenTape (Agilent Technologies, Santa Clara, CA, USA). Post-verification, the library was loaded (mixed with 10% PhiX) and sequenced with an Illumina MiSeq sequencer using a MiSeq Reagent Kit v3 (600 cycle) (Illumina Inc., San Diego, CA, USA). Following sequencing, all sub-libraries were demultiplexed to the sample level with a bespoke Python script. Tapirs, an open-source pipeline for analysing DNA metabarcoding data (https://github.com/EvoHull/Tapirs), was then utilised for taxonomic assigning demultiplexed reads. Sequence reads were quality trimmed, merged and clustered prior to taxonomic assignment against a curated UK vertebrate reference database (Harper et al., 2019). Our taxonomic assignment used a lowest common ancestor (LCA) approach based on basic local alignment search tool (BLAST) matches. Minimum identity thresholds were set to 98%.

### Inhibition

Inhibited metabarcoding samples were identified on PCR1 gels as samples with no visible target or primer dimer band and, after sequencing, samples that had a total number of assigned reads less than 500. In R1 (1 sample collected per site), inhibited samples were initially not treated differently when sequenced, resulting in 40 samples totalling less than 500 reads. As a result, 1 in 10 dilutions were later trialled on the same inhibited R1 samples and they were re-sequenced. Diluted and non-diluted R1 samples were compared against each other and the sample with more fish species detected was selected. If the number of fish species was the same, the sample with a greater number of reads was chosen. In R2, no samples were inhibited. In R3, no samples were diluted after gel inspections, but three samples totalled less than 500 assigned reads after sequencing. In R4, nine samples were diluted before sequencing after they were deemed inhibited from gels. All the R4 diluted samples totalled over assigned 500 reads when sequenced but two additional samples that were not diluted had below 500 assigned reads. Inhibition risk for qPCR was considered minimal as the master mix used in the assay is designed for use with environmental samples and is therefore much more robust to inhibition than the Q5 High-Fidelity DNA Polymerase used in the metabarcoding workflow. Additionally, the use of additional ethanol wash steps in the DNA extraction protocol after R1 largely mitigated for inhibition in metabarcoding.

### qPCR

The qPCR analysis was retrospectively performed on two targeted sub-sets of samples after all water sampling and metabarcoding sequencing had been completed:

1. all initial (single) R1 water samples (n = 145), and
2. all samples collected from *A. anguilla*-positive sites using metabarcoding (at any point during the study) (n = 132).

Given site re-sampling was based on the removal survey design using metabarcoding results, i.e. a site was not resampled when *A. anguilla* were detected with metabarcoding, it was not possible to use metabarcoding to quantify excess qPCR effort. More specially, it was not possible to calculate the number of additional qPCR samples required to detect *A. anguilla* at sites that were positive with metabarcoding and negative with PCR within the samples collected. Notwithstanding, because two samples were taken during R2-R4, one site was *A. anguilla*-positive with qPCR one sample later than the first positive metabarcoding sample. This qPCR positive sample was excluded from analysis of excess metabarcoding effort for consistency. It is also important to note that the study is a real-world comparison of two established *A. anguilla* eDNA protocols with three pooled PCR replicates for metabarcoding (Griffiths et al., 2023) and six qPCR replicates (Weldon et al., 2020), rather than an in-depth comparison of replication’s effect on detection, *per se*.

### *A. anguilla*-specific reagents

Species-specific primers and probe targeting the cytochrome B genome of *A. anguilla* were used as described in Weldon et al. (2020). The forward primer sequence was 5 ′ - TTGCCCTATTCTACCCGAACC-3′, the reverse primer sequence was 5′-ACAAGGCTAATACCCCGCC-3′, and the TaqMan probe sequence was 5′-TTGGAGACCCAGACAACTTCACCCCGGCA-3′. A synthetic gBlock standard corresponding to a 998 bp fragment of the *A. anguilla* cytochrome b gene was used as a quantitative standard.

### qPCR assay setup

Quantitative PCR assays were performed in 96-well plates. Each plate contained 12 environmental DNA samples, seven 10-fold serial dilutions of the synthetic gBlock standard (ranging from 5.77 × 10° to 5.77 × 10^6^ copies), and one no-template control (molecular-grade water). Standards and negative controls were each run in triplicate, while environmental samples were run in six replicates.

Each reaction was performed in a total volume of 15 μL, consisting of 7.5 μL TaqMan Environmental Master Mix 2.0 (Applied Biosystems), 4.75 μL molecular-grade water, 0.15 μL TaqMan probe (100 nM), 0.3 μL forward primer (200 nM), 0.3 μL reverse primer (200 nM), and 2 μL of DNA template (environmental DNA extract, standard, or molecular-grade water). All experiments were conducted using sterile equipment under laminar flow hoods. Equipment and workspaces were UV-sterilized for a minimum of 20 minutes prior to use to minimize risk of contamination.

### Cycling conditions

qPCR reactions were conducted on an Applied Biosystems StepOne Real-Time PCR System with the following thermal profile: initial incubation at 50 °C for 10 min, denaturation at 95 °C for 10 min, followed by 50 cycles of denaturation at 95 °C for 15 s and annealing/extension at 59 °C for 1 min. Plates were sealed with adhesive film to prevent cross-contamination during amplification.

### Data Analysis and method comparison

Thresholds of 20 reads were applied to *A. anguilla* for metabarcoding detection to account for low abundance of a rare target species. Thresholds 0.1% of total reads in a sample (after removal of unassigned reads) were applied to all other species. For qPCR, a single replicate amplifying was considered a positive (Weldon et al., 2020).

All statistics, analyses and data visualisation was carried out using R v4.5.1 (R Core Team, 2025) with the following packages: tidyverse (Wickham et al., 2019), VennDiagram (Chen & Boutros, 2022), viridis (Garnier et al., 2024), patchwork (Pedersen, 2025), ggpattern (Davis, 2025), vegan (Oksanen et al., 2025), iNEXT (Hsieh et al., 2020) and irr (Gamer et al., 2019). Cohen’s kappa coefficient, as used by Harper et al. (2018), was used to quantify sample-level *A. anguilla* detection agreement between qPCR and metabarcoding. The Chi-Square test of independence, also used by Harper et al. (2018), was used to test for association between qPCR and metabarcoding detections. Fisher’s exact test was used for increased confidence in the results of the chi-square test. Spearman’s rank tests were used to test for correlations between qPCR copy numbers and eDNA sequence reads for *A. anguilla*. Fisher’s exact tests were also used to test for presence absence associations between *A. anguilla* and all other fish species detected at five or more sites. A Mann-Whitney U test was used to compare fish community richness between sites where *A. anguilla* were and were not detected. PERMANOVA (Jaccard dissimilarity) was used to compare fish communities at sites with and without *A. anguilla* detections.

To assess wider metabarcoding benefits, fish species richness at sites with two or more samples was estimated using incidence-based interpolation and extrapolation in the iNEXT R package. Richness was extrapolated up to 15 samples, and 40 interpolation knots were used. Our assessment of conservation priority fish, mammal and amphibian species selected those afforded legal or policy protection under either UK or EU legislation such as the Wildlife and Countryside Act, the EC Habitats Directive, the Bern Convention and the UK Post-2010 Biodiversity Framework (see Supplementary Material). All non-native fish species in the UK were reported, whereas ubiquitous non-native mammal species, such as rat (*Rattus rattus, Rattus norvegicus*) and rabbit (*Oryctolagus cuniculus*), were not reported.

ChatGPT (OpenAI) was used to assist with generating and troubleshooting code for statistical analyses and for minor language editing to improve clarity. All outputs were reviewed, edited, and verified by the authors.

## Results

### Sample-level *A. anguilla* detection rate agreement between qPCR and metabarcoding

#### Initial (single) water sample

Of the 145 sites studied with an initial (single) water sample, 19 were positive for *A. anguilla* and qPCR was more sensitive than metabarcoding; ten (52.6%) with metabarcoding and seventeen (89.5%) with qPCR (**Error! Reference source not found**.). Eight (42.1%) were positive for *A. anguilla* with both metabarcoding and qPCR, with nine (47.4%) uniquely positive with qPCR and two (10.5%) uniquely positive with metabarcoding (Figure 1). All other initial water samples (*n* = 126) were negative with both methods.

**Figure 1.**
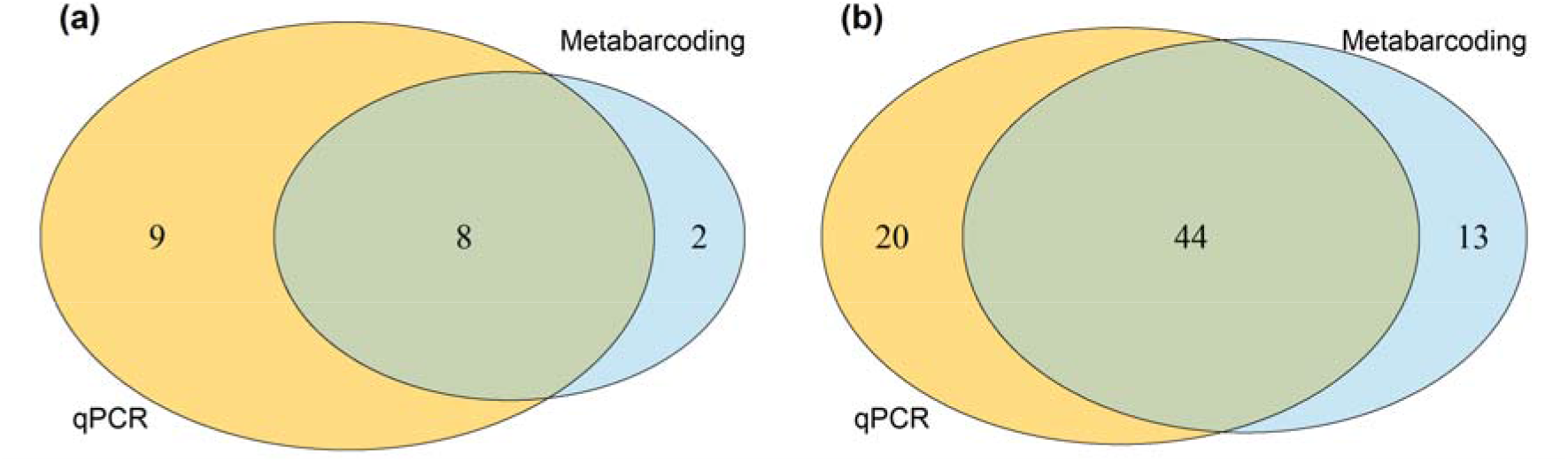
Agreement (green) in *A. anguilla* detections between qPCR (yellow) and metabarcoding (blue) for (a) the initial (single) water sample collected at each site. Samples in which *A. anguilla* was not detected by either method are not shown (n = 126). (b) for all samples analysed with both methods across the study, including the initial (single) water samples at all sites and all samples from *A. anguilla*-positive sites using metabarcoding. Samples with no detection by either method are not shown (n = 200).

### All water samples

When comparing both the initial samples from all sites and all samples from *A. anguilla*-positive sites using metabarcoding, 77 were positive for *A. anguilla*; 57 (74.0%) with metabarcoding and 64 (83.1%) with qPCR (**Error! Reference source not found**.b). Forty-four (57.1%) were positive with both methods, 20 (26.0%) were uniquely positive with qPCR and 13 (16.9%) were unique to metabarcoding (Figure 1). All other samples (n = 200) were negative with both methods.

A Spearman’s rank test found significant weak correlations between qPCR copy numbers and metabarcoding reads for *A. anguilla* (n=186, s = 776379, rho = 0.276, p<0.001), proportional reads for *A. anguilla* (n=186, s = 879742, rho = 0.180, p = 0.014) and proportional reads for *A. anguilla* with unassigned reads removed (n=186, s = 881962, rho = 0.1776156, p <0.001). A Pearson’s chi-square test with a Yates’ continuity correction (X-squared = 114.38, df = 1, p < 0.001) and a Fisher’s Exact test (*n* = 277, p <0.001) showed a significant association between the two methods. Additionally, a Cohen’s Kappa Coefficient (n = 277, z = 10.9, Kappa = 0.651, p <0.001) showed a significant moderate level of agreement between the two methods.

### Site-level *A. anguilla* detection agreement between qPCR and metabarcoding

#### *A. anguilla* detection during resampling

Of the 135 sites that were found to be negative for *A. anguilla* with an initial (single) water sample, 33 were found to be positive for *A. anguilla* with metabarcoding in later samples. For example, a further eight sites were *A. anguilla* positive with metabarcoding in the second sample, the highest number of newly positive sites were in the fourth sample (*n* = 16) and two sites had seven samples taken before *A. anguilla* presence was detected with metabarcoding (Figure 2). Only one site that was *A. anguilla*-positive with qPCR with the initial (single) water sample was consistently *A. anguilla*-negative with metabarcoding during subsequent sampling (eight samples). By contrast, *A. anguilla* were not detected by qPCR within the same number of metabarcoding samples (range = 1-7) at ten sites (total = 38 samples; Figure 2).

**Figure 2.**
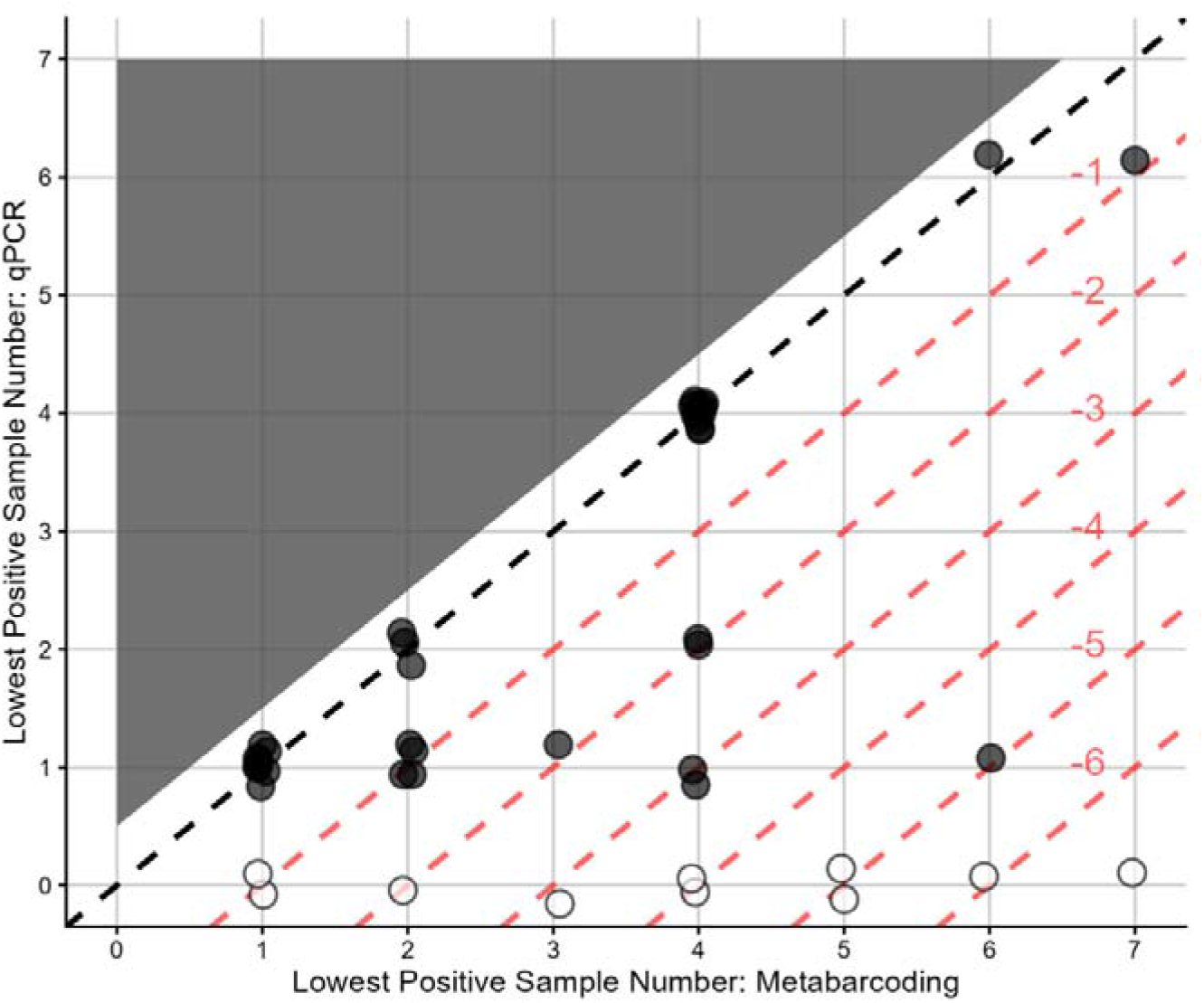
Scatter plot comparing *A. anguilla*-positive metabarcoding and qPCR sample numbers for sites where *A. anguilla* were detected using metabarcoding. The dashed black line indicates sample detection equilibrium, with points below the line representing sites where A. anguilla were detected in an earlier sample with qPCR, except “0” values (white circles) which are sites where qPCR did not detect *A. anguilla* within the same number of samples as metabarcoding.

#### Sampling effort at *A. anguilla*-positive sites

When considering excess metabarcoding effort, 33 sites were *A. anguilla*-positive with both methods and one third were *A. anguilla*-positive with qPCR in an earlier sample. Specifically, 22 (66.7%) were *A. anguilla*-positive in the same sample, five (15.2%) were one qPCR sample prior, three (9.1%) were two qPCR samples prior, two (6.1%) were three qPCR samples prior and one (3.0%) was five qPCR samples prior (Figure 2). At sites that were positive with both methods, 100 and 78 samples were required with metabarcoding and qPCR, respectively, i.e. a further 22 metabarcoding samples were required to achieve the same result as qPCR. Overall, for samples where both methods were applied, metabarcoding detected *A. anguilla* at 43 sites (with ten unique sites) using 277 samples whereas qPCR detected *A. anguilla* at 34 sites (with one unique site) using 255 samples, i.e., 26.5% more *A. anguilla*-positive sites for 8.6% more sampling effort.

### Wider metabarcoding benefits

#### Community composition

Although *A. anguilla* were the target species, metabarcoding also detected a further 28 fish species across all sites, and highest species richness at any site was 18. Fish species richness increased with additional metabarcoding samples, although the gains diminished. Specifically, across all sites, median interpolated and extrapolated fish species richness were 5.1, 8.4, 9.4 and 9.9 after one, five, ten and fifteen samples, respectively (Figure 3). In total, 615 metabarcoding samples were taken from 102 *A. anguilla*-negative sites, which culminated in 698 novel detections across 26 non-target fish species. In the eleven instances where *A. anguilla* were detected in an earlier sample with qPCR than metabarcoding, the median difference in species richness (excluding *A. anguilla*) between these samples with metabarcoding was 2, with 0 and 6 being the smallest and largest differences. It is also noteworthy that one fish (excluding *A. anguilla*), four mammal and one amphibian conservation priority species, were also detected across all sites, with *Arvicola amphibius* (n = 124) and *Cobitis taenia* (*n* = 83) being the most numerous (Figure 4). In addition, six fish and four mammal non-native species were detected, with bitterling (*Rhodeus amarus*) being the most numerous (Figure 4).

**Figure 3.**
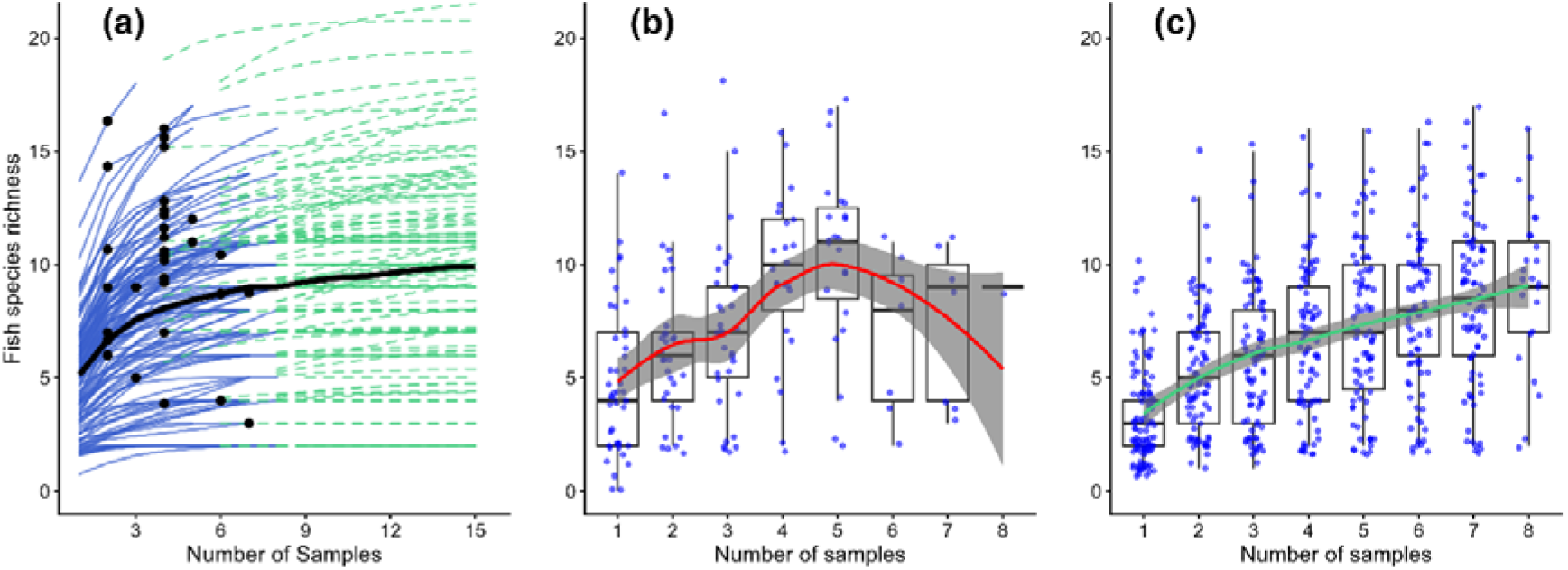
a) Fish species richness accumulation curves for all sites with metabarcoding. *A. anguilla* positive samples (black dots) and median richness (black line) detailed. Interpolated (blue line) and extrapolated (green line) data are shown. (b,c) Boxplots of median fish species richness at *A. anguilla*-positive (b) and *A. anguilla*-negative (c) sites according to the number of metabarcoding samples taken, with a red (b)/green (c) curve representing a LOESS smoothed trend with 95% confidence shading.

**Figure 4.**
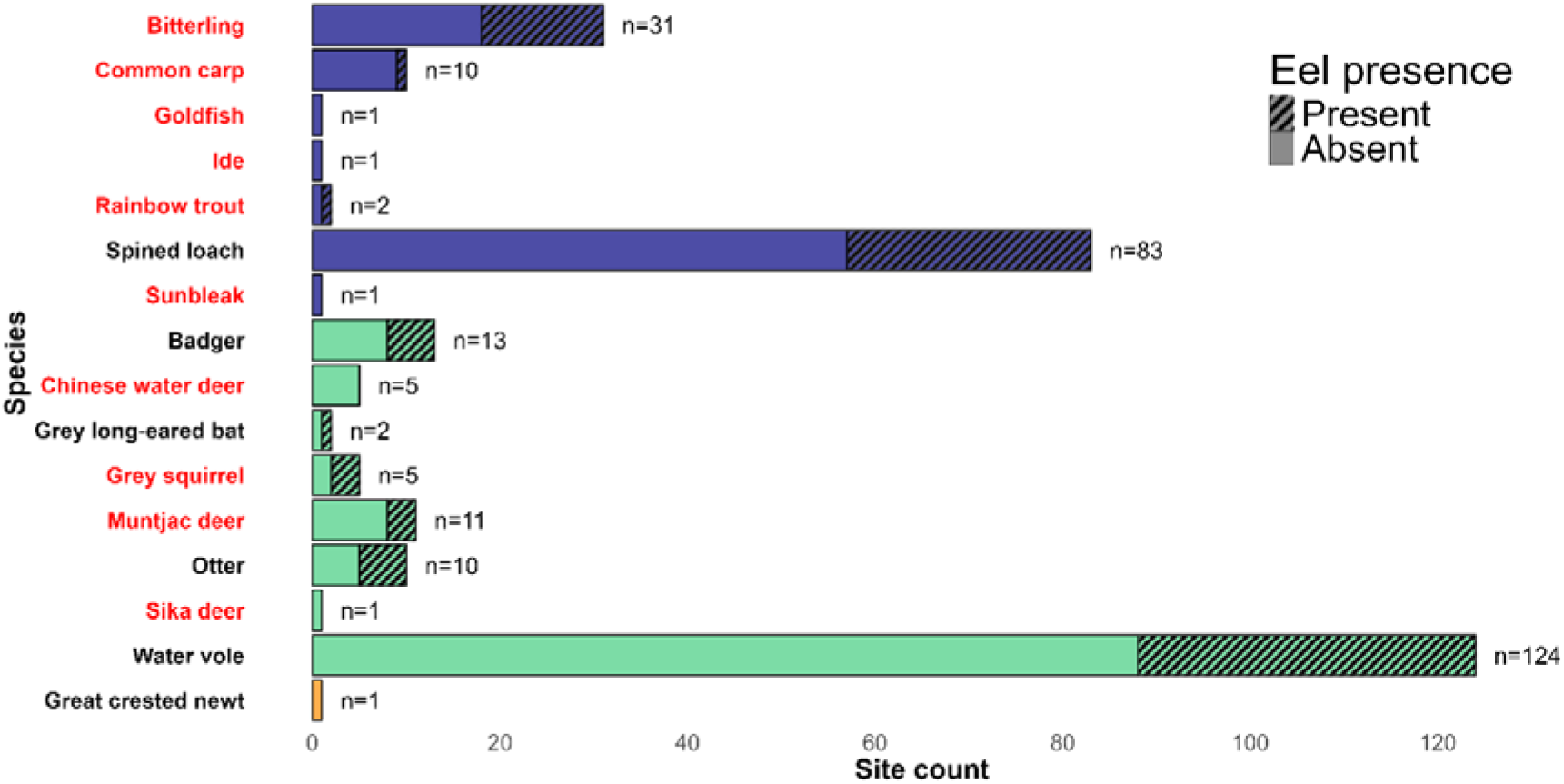
Number of sites where fish (blue bar), mammal (green bar) and amphibian (orange bar) species detected that are of interest due to conservation priority status and/or protective legislation in the UK or EU (black) or non-native in the UK (red). Bars are split by *A. anguilla* presence at sites. Details of the species’ conservation status can be found in supplementary material.

#### Species associations with *A. anguilla*

Six of the nineteen fish species detected at five or more sites were detected significantly more frequently at sites where *A. anguilla* were present, namely roach (*Rutilus rutilus*; Fisher’s exact test: OR = 4.11, p = 0.00186), dace (*Leuciscus leuciscus*; OR = 9.55, p = 0.00299), ruffe (*Gymnocephalus cernua*; OR = 2.93, p = 0.00734), common bream (*Abramis brama;* OR = 2.53, p = 0.0162), silver bream (*Blicca bjoerkna;* ; OR = 2.23, p = 0.0434) and pike (*Esox lucius*; OR = 2.69, p = 0.0317). In contrast, there were no clear associations between eel occurrence and several other species, including spined loach (*Cobitis taenia*; OR = 1.21, p = 0.714), tench (*Tinca tinca*; OR = 0.873, p = 0.718), rudd (*Scardinius erythrophthalmus*; OR = 0.934, p = 0.855) and three-spined stickleback (*Gasterosteus aculeatus*; OR = 1.57, p = 0.315). Bitterling (*Rhodeus amarus*) showed a positive but non-significant association with eel presence (OR = 2.01, p = 0.120), whereas nine-spined stickleback (*Pungitius pungitius*) tended to occur less frequently at eel sites (OR = 0.255, p = 0.0650), although this effect was not statistically significant (full details of tests for associations are in supplementary material). Interpolated fish species richness after three samples (excluding *A. anguilla*) was higher at sites where *A. anguilla* were detected (Mann-Whitney U test, W = 1946, p <0.001). No significant differences in fish community composition between *A. anguilla* positive and negative sites were found (PERMANOVA, Jaccard distance, F = 1.92, R^2^ = 0.0166, p = 0.069).

## Discussion

Across the 145 pumped catchments sampled, *A. anguilla* sample-level detection rate was higher for qPCR than metabarcoding. Specifically, 17 (89.5%) of the first water samples collected from all sites and 64 (83.1%) of all samples in the study were *A*. anguilla-positive with qPCR compared to 10 (52.6%) and 57 (74.0%) for metabarcoding, respectively. However, at a site-level, metabarcoding confirmed *A. anguilla* presence at all but one of the 34 *A. anguilla*-positive qPCR sites. At sites that were positive with both methods, there was relatively little excess metabarcoding effort with an additional 22 water samples (100 in total, compared to 78 with qPCR) to achieve the same result as qPCR. Furthermore, metabarcoding also detected ten *A. anguilla*-positive sites not found with qPCR (38 water samples). Metabarcoding also revealed a broader spectrum of vertebrate biodiversity, including 28 non-target fish species and multiple conservation priority and non-native species spanning fish, mammals and one amphibian, demonstrating its well-known wider biodiversity monitoring benefits.

The higher sample-level detection rate of qPCR than metabarcoding is consistent with prior studies that report increased sensitivity to a target species’ (Harper et al., 2018; Yu et al., 2022). It is important to note that qPCR did not detect *A. anguilla* in every positive metabarcoding sample. Thirteen *A. anguilla*-positive metabarcoding samples were missed by qPCR and therefore absolute confidence in the higher sensitivity of qPCR for any given sample cannot be assumed. While the general consensus is that qPCR offers superior sensitivity to metabarcoding, cases of imperfect detection with both methods are well documented (McCarthy et al., 2022; Johnson et al., 2024). As a result, perfect detection cannot be assumed with either method, with potential implications for method selection. A possible contributing factor to the positive metabarcoding samples missed by qPCR could be the inherent stochasticity of PCR reactions (Kelly et al., 2019). Inhibition could also be a contributor but to truly understand the effects of inhibition on both methods, a more controlled experiment using internal positive controls would be required (Opel et al., 2010).

At a site-level, when *A. anguilla* were detected with both methods, there were 22 samples of excess metabarcoding effort (100 samples) compared to qPCR (78 samples). However, this study was a retrospective comparison using samples from the removal sampling strategy using metabarcoding, and thus it was not possible to reverse this comparison to quantify excess qPCR effort, i.e. if *A. anguilla* were detected in an earlier sample with metabarcoding than qPCR. Nonetheless, had qPCR been employed earlier at these sites, only a modest amount time, money and laboratory waste – which is an increasingly relevant topic in scientific research (Alves et al., 2021; Guillardín & MacKay, 2023) – could have been saved. Ultimately, only one *A. anguilla*-positive site with qPCR in the first sample was not positive for *A. anguilla* in a later metabarcoding sample. *A. anguilla* were also not detected by qPCR in the same number or fewer samples at a further ten sites that were positive with metabarcoding. These findings are consistent with a previous study that demonstrated repeated metabarcoding sampling as an effective method for assessing fish species presence/absence with a high level of confidence (Griffiths et al., 2025b). Therefore, despite qPCR having a higher sample-level sensitivity, repeated metabarcoding sampling achieved a more robust assessments of *A. anguilla* distribution at a site-level with relatively little additional sampling effort.

Metabarcoding has the advantage of the large community data that it generates (Miya et al., 2015; Hänfling et al., 2016; Lawson Handley et al., 2019; Li et al., 2019; Griffiths et al., 2020), and fish species richness increased with additional metabarcoding sampling during this study. Given that only a relatively small amount of additional metabarcoding sampling largely mitigated its lower sample-level sensitivity compared to qPCR, it can be argued that the additional community data provided at *A. anguilla*-positive sites is worth the additional costs and effort associated with additional metabarcoding sampling. Our results were similar to other studies (Hänfling et al., 2016; Sellers et al., 2024) in showing that multiple samples are required for robust fish community composition data at sites. Furthermore, *A. anguilla* were not detected at over 70% of sites and thus metabarcoding at least provided an understanding of the site-by-site and regional fish community which would not have been provided with qPCR. The potential uses of this data are vast in both a general biodiversity context and when considering the conservation of *A. anguilla*. For example, wider fish community data can be used as indicators for assessing ecosystem health and habitat quality (Blabolil et al., 2017; Song et al., 2025) and as potential indicators of connectivity in fragmented catchments (Sun et al., 2022). Aside from *A. anguilla* conservation, numerous fish, mammals and amphibians of scientific and conservation interest were found with metabarcoding that would have been missed by qPCR, including *A. amphibius* and *C. taenia*, which are of high priority within this study region.

### Future Research

For the purpose of method comparisons, droplet digital PCR (ddPCR) and digital PCR (dPCR) could also be incorporated into further study either alongside or instead of qPCR. Other studies have already begun to include digital into eDNA methodological comparisons (Doi et al., 2015; Wood et al., 2019) and ddPCR assays have already been developed for other *Anguilla* species (Thomson-Laing et al., 2021). Multiplex qPCR/ddPCR could also be an option to provide additional community data alongside target species presence/absence (Tsuji et al., 2018; Hulley et al., 2019). However, multiplex assays are most effective when used for a small number of species and assays must be designed for each target species (Tsuji et al., 2018) whereas metabarcoding can target very large numbers of species.

## Conclusions

This regional-scale investigation of European *A. anguilla* distribution in an understudied habitat uniquely informs the debate surrounding which eDNA analysis method to use when looking to survey large areas as efficiently and effectively as possible. Although it is widely accepted that qPCR has a higher sample-level sensitivity to target taxa DNA than metabarcoding, it was only marginally the case here, with qPCR and metabarcoding detection rates being 83.1% and 74.0%, respectively. At a site-level, repeated metabarcoding sampling confirmed *A. anguilla* presence at all but one of the 34 *A. anguilla*-positive qPCR sites and *A. anguilla* were not detected by qPCR within the same number of metabarcoding samples at ten sites. At *A. anguilla*-positive sites with both methods, the amount of excess metabarcoding effort relative to qPCR was relatively small, especially when weighed against the wider metabarcoding benefits. During this study, these encompassed understanding the whole fish community, including conservation and non-native fish, amphibian and mammal species, as well as fish species associations with *A. anguilla*. With these results in mind, it is concluded that wide-scale distribution of rare and elusive species can be established with high levels of confidence using eDNA metabarcoding, without significantly compromising sensitivity, sustainability or financial efficiency.

## Acknowledgements

We are grateful to the Environment Agency for funding the study and the site owners for granting us access. Additionally, we thank R. Donnelly, S. Peixoto, C. Collins, C. Cowgill and O. Evans for assistance with sample collection and lab work. We thank D. Pollard for allowing us access to Denver Sluice Complex.

## Author Contributions

A.I.T.M carried out sample collection, lab work, data analysis and manuscript preparation. G.S.S carried out lab work, bioinformatics and data analysis. N.P.G developed the study and carried out lab work. L.L.H developed the study. B.H & R.M.W developed the study and acquired funding. J.A.M carried out lab work. J.D.B coordinated fieldwork, carried out sample collection, developed the study and acquired funding. All authors supported with manuscript preparation.

## Data availability statement

All scripts and corresponding data have been archived and made available at Zenodo: https://doi.org/10.5281/zenodo.19099273

## Ethical Statement

No live animals were handled or used in this study. Permissions were obtained from site owners and the Environment Agency prior to eDNA sampling. The eDNA sampling protocol was approved by the University of Hull ethics committee. The authors declare no conflict of interest or direct financial benefit from this study.

## References

Alves, J., Sargison, F.A., Stawarz, H., Fox, W.B., Huete, S.G., Hassan, A., McTeir, B. & Pickering, A.C. (2021) A case report: insights into reducing plastic waste in a microbiology laboratory. Access Microbiology, 3(3). Available online: 10.1099/acmi.0.000173.

Blabolil, P., Říha, M., Ricard, D., Peterka, J., Prchalová, M., Vašek, M., Čech, M., Frouzová, J., Jůza, T., Muška, M., Tušer, M., Draštík, V., Sajdlová, Z., Šmejkal, M., Vejřík, L., Matěna, J., Boukal, D.S., Ritterbusch, D. & Kubečka, J. (2017) A simple fish-based approach to assess the ecological quality of freshwater reservoirs in Central Europe. Knowledge & Management of Aquatic Ecosystems, (418), p. 53. Available online: 10.1051/kmae/2017043.

Blackman, R.C., Ling, K.K.S., Harper, L.R., Shum, P., Hänfling, B. & Lawson-Handley, L. (2020) Targeted and passive environmental DNA approaches outperform established methods for detection of quagga mussels, Dreissena rostriformis bugensis in flowing water. Ecology and Evolution, 10(23), p. 13248–13259. Available online: 10.1002/ece3.6921.

Bolland, J.D., Murphy, L.A., Stanford, R.J., Angelopoulos, N.V., Baker, N.J., Wright, R.M., Reeds, J.D. & Cowx, I.G. (2019) Direct and indirect impacts of pumping station operation on downstream migration of critically endangered European eel. Fisheries Management and Ecology, 26(1), p. 76–85. Available online: 10.1111/fme.12312.

Buysse, D., Mouton, A.M., Stevens, M., Van den Neucker, T. & Coeck, J. (2014) Mortality of European eel after downstream migration through two types of pumping stations. Fisheries Management and Ecology, 21(1), p. 13–21. Available online: 10.1111/fme.12046.

Chandelier, A., Hulin, J., Martin, G.S., Debode, F. & Massart, S. (2021) Comparison of qPCR and metabarcoding methods as tools for the detection of airborne inoculum of forest fungal pathogens. Phytopathology, 111(3), p. 570–581. Available online: 10.1094/PHYTO-02-20-0034-R.

Chen, H. & Boutros, P. (2022) VennDiagram: Generate High-Resolution Venn and Euler Plots. https://cran.r-project.org/web/packages/VennDiagram/index.html [Accessed 20 Jan 2026].

Cowgill, C., Gilbert, J.D.J., Convery, I. & Lawson Handley, L. (2025) Monitoring terrestrial rewilding with environmental DNA metabarcoding: a systematic review of current trends and recommendations. Frontiers in Conservation Science, 5. Available online: 10.3389/fcosc.2024.1473957.

Davis, T.L. (2025) Ggpattern: ‘ggplot2’ Pattern Geoms. https://cran.r-project.org/web/packages/ggpattern/index.html [Accessed 20 Jan 2026].

Doi, H., Takahara, T., Minamoto, T., Matsuhashi, S., Uchii, K. & Yamanaka, H. (2015) Droplet digital polymerase chain reaction (PCR) outperforms real-time PCR in the detection of environmental DNA from an invasive fish species. Environmental Science and Technology, 49(9), p. 5601–5608. Available online: 10.1021/acs.est.5b00253.

Evans, O.J., Norman, J., Carter, L.J., Hutchinson, T., Don, A., Wright, R.M., Tuhtan, J.A., Toming, G. & Bolland, J.D. (2024) Rethinking fish-friendliness of pumps by shifting focus to both safe and timely fish passage for effective conservation. Scientific Reports, 14(1), p. 17888. Available online: 10.1038/s41598-024-67870-5.

Gamer, M., Lemon, J., Fellows, I. & Singh, P. (2019) Irr: Various Coefficients of Interrater Reliability and Agreement. https://cran.r-project.org/web/packages/irr/index.html [Accessed 20 Jan 2026].

Garnier, S., Ross, N., Rudis, B., Filipovic-Pierucci, A., Galili, T., O’Callaghan, A., Greenwell, B., Sievert, C., Harris, D.J., Sciaini, M. & Chen, J.J. (2024) Viridis - Colorblind-Friendly Color Maps for R. https://zenodo.org/records/7890878 [Accessed 20 Jan 2026].

van Ginneken, V.J.T. & Maes, G.E. (2005) The European eel (Anguilla anguilla, Linnaeus), its lifecycle, evolution and reproduction: A literature review. Reviews in Fish Biology and Fisheries, 15(4), p. 367–398. Available online: 10.1007/s11160-006-0005-8.

Griffiths, N.P., Bolland, J.D., Wright, R.M., Blabolil, P., Macarthur, J.A., Sellers, G.S. & Hänfling, B. (2025a) Seasonal Changes in Fish eDNA Signal Vary Between Contrasting River Types. Environmental DNA, 7(3), p. e70060. Available online: 10.1002/edn3.70060.

Griffiths, N.P., Bolland, J.D., Wright, R.M., Murphy, L.A., Donnelly, R.K., Watson, H.V. & Hänfling, B. (2020) Environmental DNA metabarcoding provides enhanced detection of the European eel Anguilla anguilla and fish community structure in pumped river catchments. Journal of Fish Biology, 97(5), p. 1375–1384. Available online: 10.1111/jfb.14497.

Griffiths, N.P., Hänfling, B., Cattaneo, M., Wright, R.M., Macarthur, J.A., Peixoto, S. & Bolland, J.D. (2025b) Proving a negative: Estimating species ‘confidence in absence for decision-making’ using environmental DNA monitoring. Journal of Applied Ecology, 62(9), p. 2409–2420. Available online: 10.1111/1365-2664.70099.

Griffiths, N.P., Wright, R.M., Hänfling, B., Bolland, J.D., Drakou, K., Sellers, G.S., Zogaris, S., Tziortzis, I., Dörflinger, G. & Vasquez, M.I. (2023) Integrating environmental DNA monitoring to inform eel (Anguilla anguilla) status in freshwaters at their easternmost range—A case study in Cyprus. Ecology and Evolution, 13(2). Available online: 10.1002/ece3.9800.

Guillardín, L. & MacKay, J.J. (2023) Comparing DNA isolation methods for forest trees: quality, plastic footprint, and time-efficiency. Plant Methods, 19(1). Available online: 10.1186/s13007-023-01086-y.

Hänfling, B., Handley, L.L., Read, D.S., Hahn, C., Li, J., Nichols, P., Blackman, R.C., Oliver, A. & Winfield, I.J. (2016) Environmental DNA metabarcoding of lake fish communities reflects long-term data from established survey methods. Molecular Ecology, 25(13), p. 3101–3119. Available online: 10.1111/mec.13660.

Harper, L.R., Lawson Handley, L., Carpenter, A.I., Ghazali, M., Di Muri, C., Macgregor, C.J., Logan, T.W., Law, A., Breithaupt, T., Read, D.S., McDevitt, A.D. & Hänfling, B. (2019) Environmental DNA (eDNA) metabarcoding of pond water as a tool to survey conservation and management priority mammals. Biological Conservation, 238. Available online: 10.1016/j.biocon.2019.108225.

Harper, L.R., Lawson Handley, L., Hahn, C., Boonham, N., Rees, H.C., Gough, K.C., Lewis, E., Adams, I.P., Brotherton, P., Phillips, S. & Hänfling, B. (2018) Needle in a haystack? A comparison of eDNA metabarcoding and targeted qPCR for detection of the great crested newt (Triturus cristatus). Ecology and Evolution, 8(12), p. 6330–6341. Available online: 10.1002/ece3.4013.

Harwood, A.J.P., Perrow, M.R., Sayer, C.D., Piper, A.T., Berridge, R.J., Patmore, I.R., Emson, D. & Cooper, G. (2022) Catchment-scale distribution, abundance, habitat use, and movements of European eel (Anguilla anguilla L.) in a small UK river: Implications for conservation management. Aquatic Conservation: Marine and Freshwater Ecosystems, 32(5), p. 797–816. Available online: 10.1002/aqc.3794.

Holman, L.E., de Bruyn, M., Creer, S., Carvalho, G., Robidart, J. & Rius, M. (2019) Detection of introduced and resident marine species using environmental DNA metabarcoding of sediment and water. Scientific Reports, 9(1). Available online: 10.1038/s41598-019-47899-7.

Hsieh, T.C., Ma, K.H. & Chao, A. (2020) iNEXT: iNterpolation and EXTrapolation for species diversity. https://sites.google.com/view/chao-lab-website/software/inext [Accessed 20 Jan 2026].

Hulley, E.N., Tharmalingam, S., Zarnke, A. & Boreham, D.R. (2019) Development and validation of probe-based multiplex real-time PCR assays for the rapid and accurate detection of freshwater fish species. PLoS ONE, 14(1). Available online: 10.1371/journal.pone.0210165.

Jerde, C.L., Mahon, A.R., Chadderton, W.L. & Lodge, D.M. (2011) ‘Sight-unseen’ detection of rare aquatic species using environmental DNA. Conservation Letters, 4(2), p. 150–157. Available online: 10.1111/j.1755-263X.2010.00158.x.

Johnson, M., Tetzlaff, S., Katz, A. & Sperry, J. (2024) Comparison of qPCR and metabarcoding for environmental DNA surveillance of a freshwater parasite. Ecology and Evolution, 14(5), p. e11382. Available online: 10.1002/ece3.11382.

van Keeken, O.A., van Hal, R., Winter, H.V., Wilkes, T. & Griffioen, A.B. (2021) Migration of silver eel, Anguilla anguilla, through three water pumping stations in The Netherlands. Fisheries Management and Ecology, 28(1), p. 76–90. Available online: 10.1111/fme.12457.

Kelly, R.P., Port, J.A., Yamahara, K.M. & Crowder, L.B. (2014) Using Environmental DNA to Census Marine Fishes in a Large Mesocosm.

Kerr, J.R., Karageorgopoulos, P. & Kemp, P.S. (2015) Efficacy of a side-mounted vertically oriented bristle pass for improving upstream passage of European eel (Anguilla anguilla) and river lamprey (Lampetra fluviatilis) at an experimental Crump weir. Ecological Engineering, 85, p. 121–131. Available online: 10.1016/j.ecoleng.2015.09.013.

Kitson, J.J.N., Hahn, C., Sands, R.J., Straw, N.A., Evans, D.M. & Lunt, D.H. (2019) Detecting host– parasitoid interactions in an invasive Lepidopteran using nested tagging DNA metabarcoding. Molecular Ecology, 28(2), p. 471–483. Available online: 10.1111/mec.14518.

Knights, B., Bark, A. & Williams, B. (2009) Management of European Eel Populations in England and in Wales: a Critical Review and Pragmatic Considerations.

Laffaille, P., Baisez, A., Rigaud, C. & Feunteun, E. (2004) Habitat preferences of different European eel size classes in a reclaimed marsh: A contribution to species and ecosystem conservation. Wetlands, 24, p. 642–651.

Lawson Handley, L., Read, D.S., Winfield, I.J., Kimbell, H., Johnson, H., Li, J., Hahn, C., Blackman, R., Wilcox, R., Donnelly, R., Szitenberg, A. & Hänfling, B. (2019) Temporal and spatial variation in distribution of fish environmental DNA in England’s largest lake. Environmental DNA, 1(1), p. 26–39. Available online: 10.1002/edn3.5.

Leese, F., Sander, M., Buchner, D., Elbrecht, V., Haase, P. & Zizka, V.M.A. (2021) Improved freshwater macroinvertebrate detection from environmental DNA through minimized nontarget amplification. Environmental DNA, 3(1), p. 261–276. Available online: 10.1002/edn3.177.

Li, J., Hatton-Ellis, T.W., Lawson Handley, L.J., Kimbell, H.S., Benucci, M., Peirson, G. & Hänfling, B. (2019) Ground-truthing of a fish-based environmental DNA metabarcoding method for assessing the quality of lakes. Journal of Applied Ecology, 56(5), p. 1232–1244. Available online: 10.1111/1365-2664.13352.

Mackenzie, D.I. & Royle, J.A. (2005) Designing occupancy studies: general advice and allocating survey effort. Journal of Applied Ecology, 42(6), p. 1105–1114. Available online: 10.1111/j.1365-2664.2005.01098.x.

McCarthy, A., Rajabi, H., McClenaghan, B., Fahner, N.A., Porter, E., Singer, G.A.C. & Hajibabaei, M. (2022) Comparative analysis of fish eDNA reveals higher sensitivity achieved through targeted sequence-based metabarcoding. Molecular Ecology Resources [Preprint]. Available online: 10.1111/1755-0998.13732.

Miya, M., Sato, Y., Fukunaga, T., Sado, T., Poulsen, J.Y., Sato, K., Minamoto, T., Yamamoto, S., Yamanaka, H., Araki, H., Kondoh, M. & Iwasaki, W. (2015) MiFish, a set of universal PCR primers for metabarcoding environmental DNA from fishes: Detection of more than 230 subtropical marine species. Royal Society Open Science, 2(7). Available online: 10.1098/rsos.150088.

Mychek-Londer, J.G., Balasingham, K.D. & Heath, D.D. (2020) Using environmental DNA metabarcoding to map invasive and native invertebrates in two Great Lakes tributaries. Environmental DNA, 2(3), p. 283–297. Available online: 10.1002/edn3.56.

Nichols, J.D. & Williams, B.K. (2006) Monitoring for conservation. Trends in Ecology & Evolution, 21(12), p. 668–673. Available online: 10.1016/j.tree.2006.08.007.

Norman, J., Reeds, J., Wright, R.M. & Bolland, J.D. (2023a) The impact of extreme flood-relief pump operations on resident fish in an artificial drain and the potential for artificial habitat introduction. Fisheries Management and Ecology [Preprint].

Norman, J., Wright, R.M., Don, A. & Bolland, J.D. (2023b) Understanding the temporal dynamics of a lowland river fish community at a hazardous intake and floodgate to inform safe operation. Journal of Environmental Management, 336. Available online: 10.1016/j.jenvman.2023.117716.

Nunn, A.D., Tewson, L.H., Bolland, J.D., Harvey, J.P. & Cowx, I.G. (2014) Temporal and spatial variations in the abundance and population structure of the spined loach (Cobitis taenia), a scarce fish species: implications for condition assessment and conservation. Aquatic Conservation: Marine and Freshwater Ecosystems, 24(6), p. 818–830. Available online: 10.1002/aqc.2451.

Oksanen, J., Simpson, G.L., Blanchet, F.G., Kindt, R., Legendre, P., Minchin, P.R., O’Hara, R.B., Solymos, P., Stevens, M.H.H., Szoecs, E., Wagner, H., Barbour, M., Bedward, M., Bolker, B., Borcard, D., Borman, T., Carvalho, G., Chirico, M., Caceres, M.D., Durand, S., Evangelista, H.B.A., FitzJohn, R., Friendly, M., Furneaux, B., Hannigan, G., Hill, M.O., Lahti, L., Martino, C., McGlinn, D., Ouellette, M.-H., Cunha, E.R., Smith, T., Stier, A., Braak, C.J.F.T. & Weedon, J. (2025) Vegan: Community Ecology Package. https://cran.r-project.org/web/packages/vegan/index.html [Accessed 20 Jan 2026].

Opel, K.L., Chung, D. & McCord, B.R. (2010) A study of PCR inhibition mechanisms using real time PCR. In. Journal of Forensic Sciences, p. 25–33. Available online: 10.1111/j.1556-4029.2009.01245.x.

Ouellet, V., Collins, M.J., Kocik, J.F., Saunders, R., Sheehan, T.F., Ogburn, M.B. & Trinko Lake, T. (2022) The diadromous watersheds-ocean continuum: Managing diadromous fish as a community for ecosystem resilience. Frontiers in Ecology and Evolution, 10. Available online: 10.3389/fevo.2022.1007599.

Pedersen, T.L. (2025) Patchwork: The Composer of Plots. https://cran.r-project.org/web/packages/patchwork/index.html [Accessed 20 Jan 2026].

Peixoto, S., Blackman, R.C., Porter, J., Wan, A., Gerrard, C., Aston, B. & Handley, L.L. (2023) Mussels on the move: new records of the invasive non-native quagga mussel (Dreissena rostriformis bugensis) in Great Britain using eDNA and a new probe-based qPCR assay. bioRxiv [Preprint]. Available online: 10.1101/2023.12.18.572119.

Penaluna, B.E., Allen, J.M., Arismendi, I., Levi, T., Garcia, T.S. & Walter, J.K. (2021) Better boundaries: identifying the upper extent of fish distributions in forested streams using eDNA and electrofishing. Ecosphere, 12(1), p. e03332. Available online: 10.1002/ecs2.3332.

Pike, C., Crook, V., Gollock, M., Beaulaton, L., Belpaire, C., Dekker, W., Díaz, E., Durif, C.M.F. & Hanel, R. (2020) Anguilla anguilla. IUCN. https://hal.science/hal-04461073/document.

Podgorniak, T., Blanchet, S., de Oliveira, E., Daverat, F. & Pierron, F. (2016) To boldly climb: Behavioural and cognitive differences in migrating European glass eels. Royal Society Open Science, 3(1). Available online: 10.1098/rsos.150665.

Pont, D., Meulenbroek, P., Bammer, V., Dejean, T., Erős, T., Jean, P., Lenhardt, M., Nagel, C., Pekarik, L., Schabuss, M., Stoeckle, B.C., Stoica, E., Zornig, H., Weigand, A. & Valentini, A. (2022) Quantitative monitoring of diverse fish communities on a large scale combining eDNA metabarcoding and qPCR. Molecular Ecology Resources [Preprint]. Available online: 10.1111/1755-0998.13715.

Quail, M.A., Swerdlow, H. & Turner, D.J. (2009) Improved Protocols for the Illumina Genome Analyzer Sequencing System. Current Protocols in Human Genetics, 62(1), p. 18.2.1-18.2.27. Available online: 10.1002/0471142905.hg1802s62.

R Core Team (2025) R: A language and environment for statistical computing. R Foundation for Statistical Computing, Vienna, Austria. https://cir.nii.ac.jp/crid/1370013168792282134 [Accessed 20 Jan 2026].

Riaz, T., Shehzad, W., Viari, A., Pompanon, F., Taberlet, P. & Coissac, E. (2011) EcoPrimers: Inference of new DNA barcode markers from whole genome sequence analysis. Nucleic Acids Research, 39(21). Available online: 10.1093/nar/gkr732.

Richards, D.R., Maltby, L., Moggridge, H.L. & Warren, P.H. (2014) European water voles in a reconnected lowland river floodplain: habitat preferences and distribution patterns following the restoration of flooding. Wetlands Ecology and Management, 22(5), p. 539–549. Available online: 10.1007/s11273-014-9350-x.

Sellers, G.S., Di Muri, C., Gómez, A. & Hänfling, B. (2018) Mu-DNA: A modular universal DNA extraction method adaptable for a wide range of sample types. Metabarcoding and Metagenomics, 2. Available online: 10.3897/mbmg.2.24556.

Sellers, G.S., Jerde, C.L., Harper, L.R., Benucci, M., Di Muri, C., Li, J., Peirson, G., Walsh, K., Hatton-Ellis, T., Duncan, W., Duguid, A., Ottewell, D., Willby, N., Law, A., Bean, C.W., Winfield, I.J., Read, D.S., Handley, L.L. & Hänfling, B. (2024) Optimising species detection probability and sampling effort in lake fish eDNA surveys. Metabarcoding and Metagenomics, 8. Available online: 10.3897/mbmg.8.104655.

Song, T., Zi, F., Huang, Y., Fang, L., Zhang, Y., Liu, Y., Chang, J. & Li, J. (2025) Assessment of Aquatic Ecosystem Health in the Irtysh River Basin Using eDNA Metabarcoding. Water, 17(2), p. 246. Available online: 10.3390/w17020246.

Sun, J., Tummers, J.S., Galib, S.M. & Lucas, M.C. (2022) Fish community and abundance response to improved connectivity and more natural hydromorphology in a post-industrial subcatchment. Science of The Total Environment, 802, p. 149720. Available online: 10.1016/j.scitotenv.2021.149720.

Thomson-Laing, G., Parai, R., Kelly, L.T., Pochon, X., Newnham, R., Vandergoes, M.J., Howarth, J.D. & Wood, S.A. (2021) Development of droplet digital Polymerase Chain Reaction assays for the detection of long-finned (Anguilla dieffenbachii) and short-finned (Anguilla australis) eels in environmental samples. PeerJ, 9. Available online: 10.7717/peerj.12157.

Tsuji, S., Iguchi, Y., Shibata, N., Teramura, I., Kitagawa, T. & Yamanaka, H. (2018) Real-time multiplex PCR for simultaneous detection of multiple species from environmental DNA: An application on two Japanese medaka species. Scientific Reports, 8(1). Available online: 10.1038/s41598-018-27434-w.

Verhelst, P., Reubens, J., Buysse, D., Goethals, P., Van Wichelen, J. & Moens, T. (2021) Toward a roadmap for diadromous fish conservation: the Big Five considerations. Frontiers in Ecology and the Environment, 19(7), p. 396–403. Available online: 10.1002/fee.2361.

Weldon, L., O’Leary, C., Steer, M., Newton, L., Macdonald, H. & Sargeant, S.L. (2020) A comparison of European eel Anguilla anguilla eDNA concentrations to fyke net catches in five Irish lakes. Environmental DNA, 2(4), p. 587–600. Available online: 10.1002/edn3.91.

White, N.E., Guzik, M.T., Austin, A.D., Moore, G.I., Humphreys, W.F., Alexander, J. & Bunce, M. (2020) Detection of the rare Australian endemic blind cave eel (Ophisternon candidum) with environmental DNA: implications for threatened species management in subterranean environments. Hydrobiologia, 847(15), p. 3201–3211. Available online: 10.1007/s10750-020-04304-z.

Wickham, H., Averick, M., Bryan, J., Chang, W., McGowan, L.D., François, R., Grolemund, G., Hayes, A., Henry, L., Hester, J., Kuhn, M., Pedersen, T.L., Miller, E., Bache, S.M., Müller, K., Ooms, J., Robinson, D., Seidel, D.P., Spinu, V., Takahashi, K., Vaughan, D., Wilke, C., Woo, K. & Yutani, H. (2019) Welcome to the Tidyverse. Journal of Open Source Software, 4(43), p. 1686. Available online: 10.21105/joss.01686.

Wilcox, T.M., McKelvey, K.S., Young, M.K., Sepulveda, A.J., Shepard, B.B., Jane, S.F., Whiteley, A.R., Lowe, W.H. & Schwartz, M.K. (2016) Understanding environmental DNA detection probabilities: A case study using a stream-dwelling char Salvelinus fontinalis. Biological Conservation, 194, p. 209–216. Available online: 10.1016/j.biocon.2015.12.023.

Wood, S.A., Pochon, X., Laroche, O., von Ammon, U., Adamson, J. & Zaiko, A. (2019) A comparison of droplet digital polymerase chain reaction (PCR), quantitative PCR and metabarcoding for species-specific detection in environmental DNA. Molecular Ecology Resources, 19(6), p. 1407–1419. Available online: 10.1111/1755-0998.13055.

Yu, Z., Ito, S.I., Wong, M.K.S., Yoshizawa, S., Inoue, J., Itoh, S., Yukami, R., Ishikawa, K., Guo, C., Ijichi, M. & Hyodo, S. (2022) Comparison of species-specific qPCR and metabarcoding methods to detect small pelagic fish distribution from open ocean environmental DNA. PLoS ONE, 17(9 September). Available online: 10.1371/journal.pone.0273670.

