## Supplementary for "Species-specific versus community-wide assays in eDNA monitoring of the European eel *Anguilla anguilla*: Trade-offs between detection sensitivity and the value of additional community data"

Table 1: Designations for Conservation priority species in Figure 4

| Common name | Latin name | Conservation status |
| --- | --- | --- |
| Spined Loach | *Cobitis taenia* | - Anex II, EC Habitats Directive - Appendix III, Bern Convention |
| Badger | *Meles meles* | - UK Protection of Badgers Act, 1992 - UK Wildlife and Countryside Act, 1981 |
| Grey long-eared bat | *Plecotus austriacus* | - UK Wildlife and Countryside Act, 1981 - Anex IV, EC Habitats Directive - IUCN: Near-threatened |
| Otter | *Lutra lutra* | - UK Wildlife and Countryside Act, 1981 - UK Post-2010 Biodiversity Framework Priority Species - Anex IV, EC Habitats Directive - IUCN: Near-threated |
| Water vole | *Arvicola amphibius* | - UK Wildlife and Countryside Act, 1981 - UK Post-2010 Biodiversity Framework Priority Species |
| Great crested newt | *Triturus cristatus* | - UK Wildlife and Countryside Act, 1981 - UK Post-2010 Biodiversity Framework Priority Species - Anex IV, EC Habitats Directive |

Table 2: fisher’s exact test results for fish associations with European eel (*Anguilla anguilla*)

| **Species** | **Odds Ratio** | **95% CI Lower** | **95% CI Upper** | **p-value** |
| --- | --- | --- | --- | --- |
| *Rutilus rutilus* | 4.108 | 1.531 | 13.001 | 0.0019 |
| *Cobitis taenia* | 1.206 | 0.552 | 2.681 | 0.7139 |
| *Gymnocephalus cernua* | 2.927 | 1.253 | 6.885 | 0.0073 |
| *Esox lucius* | 2.691 | 1.082 | 7.424 | 0.0317 |
| *Perca fluviatilis/Sander lucioperca* | 2.027 | 0.875 | 4.978 | 0.0904 |
| *Squalius cephalus* | 0.947 | 0.087 | 6.074 | 1 |
| *Gobio gobio* | 2.325 | 0.962 | 5.59 | 0.0522 |
| *Leuciscus leuciscus* | 9.546 | 1.714 | 98.342 | 0.003 |
| *Barbatula barbatula* | 2.097 | 0.856 | 5.08 | 0.0836 |
| *Scardinius erythrophthalmus* | 0.934 | 0.427 | 2.066 | 0.8554 |
| *Abramis brama* | 2.529 | 1.154 | 5.649 | 0.0162 |
| *Pungitius pungitius* | 0.255 | 0.05 | 1.142 | 0.065 |
| *Gasterosteus aculeatus* | 1.569 | 0.636 | 4.186 | 0.315 |
| *Tinca tinca* | 0.873 | 0.402 | 1.907 | 0.7183 |
| *Rhodeus amarus* | 2.012 | 0.803 | 4.964 | 0.1201 |
| *Cyprinus carpio* | 0.248 | 0.005 | 1.89 | 0.2817 |
| *Blicca bjoerkna* | 2.232 | 0.963 | 5.161 | 0.0434 |
| *Alburnus alburnus* | 1.604 | 0.13 | 14.543 | 0.6332 |
| *Cottus gobio* | 2.187 | 0.567 | 8.176 | 0.2061 |
